# Pupil dynamics-derived sleep stage classification of a head-fixed mouse using a recurrent neural network

**DOI:** 10.1101/2022.08.06.503067

**Authors:** Goh Kobayashi, Kenji F. Tanaka, Norio Takata

## Abstract

The standard method for sleep state classification is thresholding amplitudes of electroencephalography (EEG) and electromyography (EMG), followed by an expert’s manual correction. Although popular, the method entails some shortcomings: 1) the time-consuming manual correction by human experts is sometimes a bottleneck hindering sleep studies; 2) EEG electrodes on the skull interfere with wide-field imaging of the cortical activity of a head-fixed mouse under a microscope; 3) invasive surgery to fix the electrodes on the thin skull of a mouse risks brain tissue injury; and 4) metal electrodes for EEG and EMG are difficult to apply to some experiment apparatus such as that for functional magnetic resonance imaging. To overcome these shortcomings, we propose a pupil dynamics-based vigilance state classification for a head-fixed mouse using a long short-term memory (LSTM) model, a variant of recurrent neural networks, for multi-class labeling of NREM, REM, and WAKE states. For supervisory hypnography, EEG and EMG recording were performed for a head-fixed mouse, combined with left eye pupillometry using a USB camera and a markerless tracking toolbox, DeepLabCut. Our open-source LSTM model with feature inputs of pupil diameter, location, velocity, and eyelid opening for 10 s at a 10 Hz sampling rate achieved vigilance state estimation with a higher classification performance (macro F1 score, 0.77; accuracy, 86%) than a feed forward neural network. Findings from diverse pupillary dynamics implied subdivision of a vigilance state defined by EEG and EMG. Pupil dynamics-based hypnography can expand the scope of alternatives for sleep stage scoring of head fixed mice.

## Introduction

Sleep stage classification is indispensable to investigate the function, mechanism, and pathology of sleep.^1–4^ Vigilance states of rodents are commonly divided into three: wake period (WAKE), rapid eye movement (REM) sleep, and non-REM (NREM) sleep.^5,6^ The standard procedure for vigilance state classification is thresholding amplitudes of electroencephalogram (EEG) and electromyogram (EMG), followed by experts’ manual correction.^7^ Although they are accepted widely and used routinely,^8^ several important shortcomings hinder EEG-based and EMG-based sleep stage classification: 1) Manual inspection of hypnograms by human experts is time-consuming and sometimes poses a bottleneck that impedes sleep studies.^9–11^ Manual scoring is nevertheless necessary because of individual differences of EEG-thresholds and EMG-thresholds among mice, and because of fluctuation of the thresholds, even in the same mouse, over time. 2) Also, EEG electrodes on the skull interfere with wide-field imaging of cortical activity,^12–16^ preventing investigation of bilateral cortical activity during sleep.^17^ 3) Invasive surgery necessary to fix an EEG electrode on the thin skull of a mouse risks brain tissue injury and infection that might modulate the structure and function of the cortex.^18,19^ 4) Applying EEG and EMG recording for functional magnetic resonance imaging (fMRI) is difficult in mice because a susceptibility artifact of the metal electrodes on the thin skull of a mouse deteriorates cortical fMRI images,^20,21^ thereby allowing fMRI only in anesthetized or awake but not sleeping mice.^22^ Earlier investigations of a vigilance state classification without EEG and EMG recording have addressed some but not all shortcomings outlined above.^23–25^

Pupil dynamics is a promising candidate for hypnogram construction without recording of EEG and EMG because eyes are closely associated with sleep.^26–28^ The association has been investigated intensively in human subjects.^29^ During wake and alert periods, pupils are large.^30^ Immediately before falling asleep, eyelids begin to droop, eyes move upward, pupils contract strongly, and the pupil diameter begins to drift erratically.^31^ During sleep, eye movements are slow and pendular. The pupil diameter shows spontaneous waves of constriction and dilation, which were observed in a human subject using a lid crutch.^30^ Rapid and jerky eye movements appear during REM sleep.^32^ Rodent studies revealed tight coupling between pupil dynamics and neuronal activities.^33–35^ Whole-cell patch-clamp recordings in awake mice have demonstrated that pupil constriction and dilation respectively accompanied synchronization and desynchronization of membrane potentials of cortical neurons.^34^ Rapid and longer-lasting pupil dilations respectively coincide with phasic and sustained activities in noradrenergic and cholinergic axons.^36^ An active role of pupillary constriction was demonstrated to stabilize deep sleep.^33^ From that study, they succeeded in estimating the vigilance states of head-fixed mice based on pupil diameter using a feed forward neural network (FFNN) with a hidden layer of 30 nodes, achieving estimation accuracies of 96% for NREM, 58% for REM, and 95% for WAKE. However, the estimation was limited to a vigilance state lasting more than 100 s, as confirmed from EEG and EMG recording. Therefore, it remains unknown whether pupil dynamics can adequately represent the moment-by-moment dynamics of vigilant states.

Neural network models have been applied for sleep stage scoring dominantly with EEG and EMG data.^37^ Deep learning models became popular in recent years, typical of which include FFNNs and recurrent neural networks (RNNs). A convolutional neural network (CNN), a variant of FFNNs, combined with a hidden Markov model (HMM), was trained on 4-s bins of EEG and EMG data from only two mice, achieving automatic sleep stage scoring with accuracy of 93–99%, with generalization capabilities.^38^ A long short-term memory (LSTM) model, a variant of RNNs, is regarded as an effective model for a sequence classification because the “forget gate” in its architecture facilitates the efficient usage of past information in a bin.^39,40^ An LSTM model combined with a CNN was trained using 20-s bins of EEG and EMG data from 4,200 mice, demonstrating sleep stage classification with accuracy of 96.6%.^11^ Although these state-of-the-art architectures on EEG and EMG data resolved the time-consuming visual inspection of a hypnogram (shortcoming 1) and achieved sleep scoring performance that was comparable to or exceeding the inter-rater agreement rate by human experts (88–95%),^11,41^ these models were not free from shortcomings 2–4 of EEG and EMG recording presented above. A recent attempt to apply machine learning to a 10-s bin video of a freely moving mouse for sleep staging instead of EEG and EMG measurement achieved accuracy of 92 ± 5% (mean ± SD).^10^ This machine vision method resolved the time-consuming manual correction of a hypnogram (shortcoming 1) and invasive surgery (shortcoming 3), but applying it for a head-fixed mouse under a microscope (shortcoming 2) or fMRI (shortcoming 4) might be difficult because the visual classification of sleep relied on the movement and posture of mice.

As described in this paper, we propose a pupil-dynamics derived hypnogram of a head-fixed mouse using a simple one-layer LSTM model. A supervisory hypnogram with 10-s bin was prepared by EEG and EMG recording from 4 male C57BL/6J mice. Simultaneous monitoring of a left eye by a USB camera in combination with a markerless tracking toolbox DeepLabCut^42^ provided input features for LSTM models on pupil diameter, location, velocity, and eyelid opening at 10 Hz. Comparison of LSTM models with distinct hyperparameters on input features and other parameters was performed with a five-fold crossvalidation technique using data from three mice. Performance of the best LSTM model was tested using the data of the reserved mouse, achieving accuracy of 86% and macro F1 of 0.77, which were higher than those of an FFNN model. Additionally, we noticed characteristic pupillary dynamics that indicated substates within a vigilance state in an EEG and EMG based hypnogram. Consequently, a pupil dynamics-derived hypnogram would add an attractive choice for sleep staging of a head-fixed mouse.

## Materials and Methods

### Ethics Statement

All animal experiments were conducted in accordance with the National Institutes of Health Guide for Care and Use of Laboratory Animals (NIH Publications No. 8023) and were approved by the Animal Ethics Committee of Keio University (approval number: 18076-(2)).

### Surgery and habituation

Four male C57BL/6 mice, eight weeks postnatal and weighing 22–27 g at the start of the experiment, were purchased from CLEA Japan, Inc. (Tokyo, Japan). The animals were deeply anesthetized with a mixture of ketamine and xylazine (100 mg/kg and 10 mg/kg, respectively, i.p.), and were fixed to a stereotaxic apparatus (SM-15; Narishige Scientific Instrument, Tokyo, Japan). After shaving the hair on their head and neck, a longitudinal incision was made on the scalp, exposing the skull surface. The periosteum and blood on the skull were removed thoroughly. Three holes (ϕ1 mm) for EEG recording were prepared using a dental drill (ϕ0.8 mm, 23017; Nakanishi Inc., Tochigi, Japan) on the skull without penetration. These holes were made to accommodate the following: 1) a reference electrode at 1.0 mm anterior to the Bregma (AP +1.0 mm) and 1.5 mm lateral from the midline to the right (ML +1.5 mm) on the frontal skull; 2) a signal electrode at 1.0 mm anterior to the Lambda and ML +2.5 mm on the parietal skull; and 3) a ground electrode at the center of the interparietal skull (ML 0 mm). EEG electrodes, consisted of a silver wire (#786500; A-M Systems Inc., WA, USA) and a screw (ϕ1 mm, length 2 mm; #0-1 nabe; Yamazaki Corp., Shiga, Japan), were screwed to the holes after attaching a metal chamber frame (CF-10; Narishige Scientific Instrument, Tokyo, Japan) on the skull using dental acrylic (Super-Bond C&B; Sun Medical, Shiga, Japan). EMG electrodes, consisting of a flexible stainless steel (AS633; Cooner Wire, CA, USA), were inserted into the trapezius muscles bilaterally. EEG and EMG wires were fixed to the skull using the dental acrylic.

For recovery after surgery, animals were single-housed with water and food pellets provided ad libitum at ambient temperature (22–24 °C) on a 12–hr light:dark cycle (light on at 8:00 h and off at 20:00 h) for at least a week. Mice were then habituated to sleep under a head-restrained condition in a dark room.^22^ The duration for the habituation increased gradually from 2 hr per day at the beginning to the maximum of 6 hr per day. The habituation was performed at least a week. At the beginning of habituation, mice were mostly wake under the head-fixed situation. During the latter part of the habituation, mice showed a characteristic polyphasic sleep under the head-fixed condition (Fig. 1C). Six-hour habituation seemed critical: repeated 2 hr habituation failed to make head-fixed mice sleep under these experimental conditions.

**Fig. 1.**
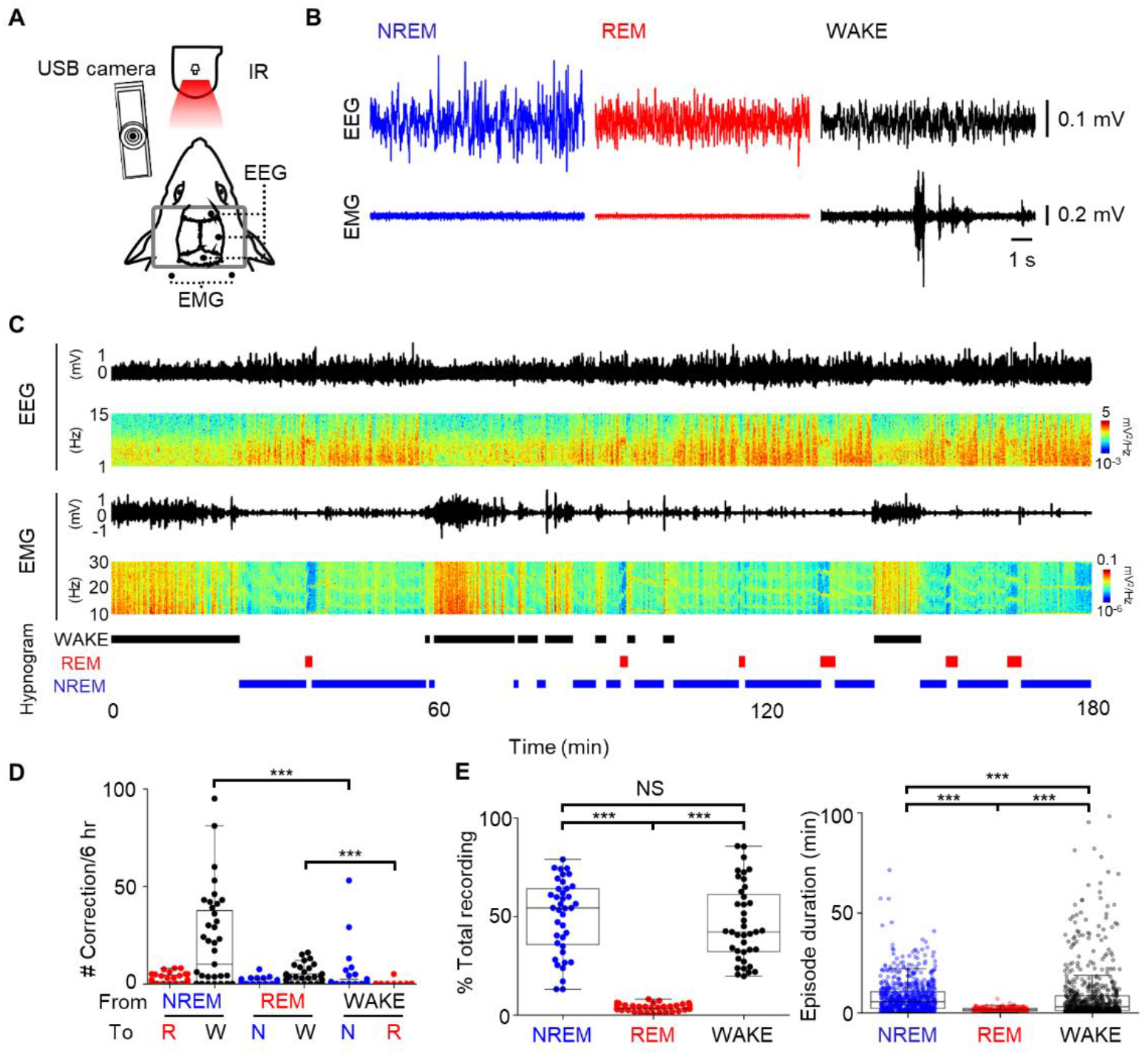
Preparation of a supervisory hypnogram using EEG and EMG as ground truth labels for deep learning. **A:** Schematic of EEG and EMG recordings and pupillometry from a head-fixed mouse. A head frame (gray rectangle) was attached to the skull for fixation. **B:** Representative 10-s EEG and EMG traces during NREM (blue), REM (red), and WAKE (black) periods. **C:** Representative EEG and EMG traces for a single recording session with a supervisory hypnogram. Blue columns in the EMG spectrogram represent a marked decrease of EMG power during REM sleep. **D:** Number of manual corrections required for the threshold-based hypnogram (Threshold) to prepare a supervisory hypnogram. Corrections from sleep states (NREM or REM) to WAKE states were significantly more numerous than vice versa (two-sided Wilcoxon signed-ranks test, *n* = 39 sessions from 4 mice. Counts of episode corrections per 6 hr: 21 ± 24 episodes from NREM to WAKE vs. 3 ± 10 episodes from WAKE to NREM; 4 ± 5 episodes from REM to WAKE vs. 0.1 ± 0.8 episodes from WAKE to REM. Data are mean ± SD, \*\*\**p* < 0.0001). **E:** Time spent in each vigilance state in a single recording session (left) and duration for an individual episode in all recording session (right) based on a supervisory hypnogram. Those values corresponded to those obtained for freely moving mice (see Results). Mean % total recording for NREM (48.9 ± 3.0%) and WAKE (47.6 ± 3.1%) were significantly greater than those for REM (3.5 ± 0.3%; *n* = 29 sessions from 4 mice. Steel-Dwass test, \*\*\**p* < 0.0001). Mean episode durations for NREM, REM, and WAKE were distinct (NREM, 8.2 ± 11.3 min; REM 1.7 ± 0.9 min; WAKE 9.1 ± 19.4 min, [mean ± SD]; Steel-Dwass test, \*\*\**p* < 0.0001).

### Electrophysiology and pupillometry

During a recording session, EEG, EMG, and left eye pupillometry were performed under a head-fixed condition that lasted 3–6 hr per day in light phase within zeitgeber time (ZT) = 2-9 (10 AM to 6 PM). Specifically, EEG and EMG were amplified 1,000 times, respectively, with filter configurations of 1–300 Hz and 10–300 Hz (Model 3000; A-M Systems Inc., WA, USA), analog-to-digital converted (16-bit depth; NI-9215; National Instruments Corp., TX, USA), and recorded at a sampling rate of 1,000 Hz using a custom written program (LabVIEW 2016; National Instruments Corp., TX, USA) on a computer (Windows 10 Pro; Microsoft Corp., WA, USA). Images of a mouse left eye were captured using a custom written program (LabVIEW 2016; National Instruments Corp., WA, USA) on the PC at 30.3 ± 16.5 Hz (mean ± SD) with an infrared (IR) LED ring light (940 nm, FRS5 JS; OptoSupply Ltd., Hong Kong) and a USB camera for which the IR cutoff filter was removed beforehand (BSW200MBK; Buffalo Inc., Japan). A single recording session was performed once a day for 3–6 hr.

### Pupil dynamics feature extraction

A deep learning-based animal pose estimation software, DeepLabCut (ver. 2.2rc3), was used to acquire position of a pupil rostral (PR), a pupil caudal (PC), upper eyelid margin (UEM), and lower eyelid margin (LEM) in the left eye images (Fig. 2A).^42,43^ We first manually extracted 600 left-eye images in all from 17 recording sessions of 4 mice (150 ± 167 images/mouse [mean ± SD]). The manual selection of eye images improved positioning performance by DeepLabCut because rare or difficult images for positioning, e.g., pupil images with overlapping whiskers or IR-light reflection, can be selected intentionally to train DeepLabCut, which was difficult or impossible to accomplish using automatic image extraction by DeepLabCut through k-means clustering or a random temporal sampling. We manually labeled the positions of the PR, PC, UEM, and LEM in the extracted images using the graphical user interface (GUI) of DeepLabCut. Specifically, we defined first a line connecting the inner and outer corner of the eye (I-O line; a white dotted line in Fig. 2A). Parallel to the I-O line, a line segment was obtained connecting the rostral and caudal edges of a pupil with the largest distance (an orange dashed line in Fig. 2B). These edges were defined respectively as positions for PR (purple dot in Fig. 2A) and PC (light blue dot in Fig. 2A). The intersections of the vertical bisector of the I-O line with upper and lower eyelids were defined as a UEM (a yellow dot in Fig. 2A) and a LEM (a red dot in Fig. 2A). If whiskers or IR-light reflection overlapped with any position, then a slightly displaced location was assigned to avoid the overlap. The 95% of the manually obtained dataset (570 from the 600 left eye images) was used to train DeepLabCut (a left panel in Fig. 2B). We used a ResNet-50-based neural network with default parameters for 1,030,000 number of training iterations.^44^ After the learning curve reached a plateau, the generalization performance of DeepLabCut was evaluated with root mean square error (RMSE) using a remaining test dataset (30 from the 600 left eye images; a right bar graph in Fig. 2B). After confirming successful positioning by DeepLabCut, the four positions in the left eye images were obtained automatically by DeepLabCut for 22,589,940 images in 39 sessions from 4 mice. The DeepLabCut was run on a local computer (MacOS 10.15 Catalina; Apple Computer Inc.) for manual extraction and labeling of eye images, and on a Google Cloud server using Google Colaboratory Pro (Alphabet Inc.) for training and automatic positioning.

**Fig. 2.**
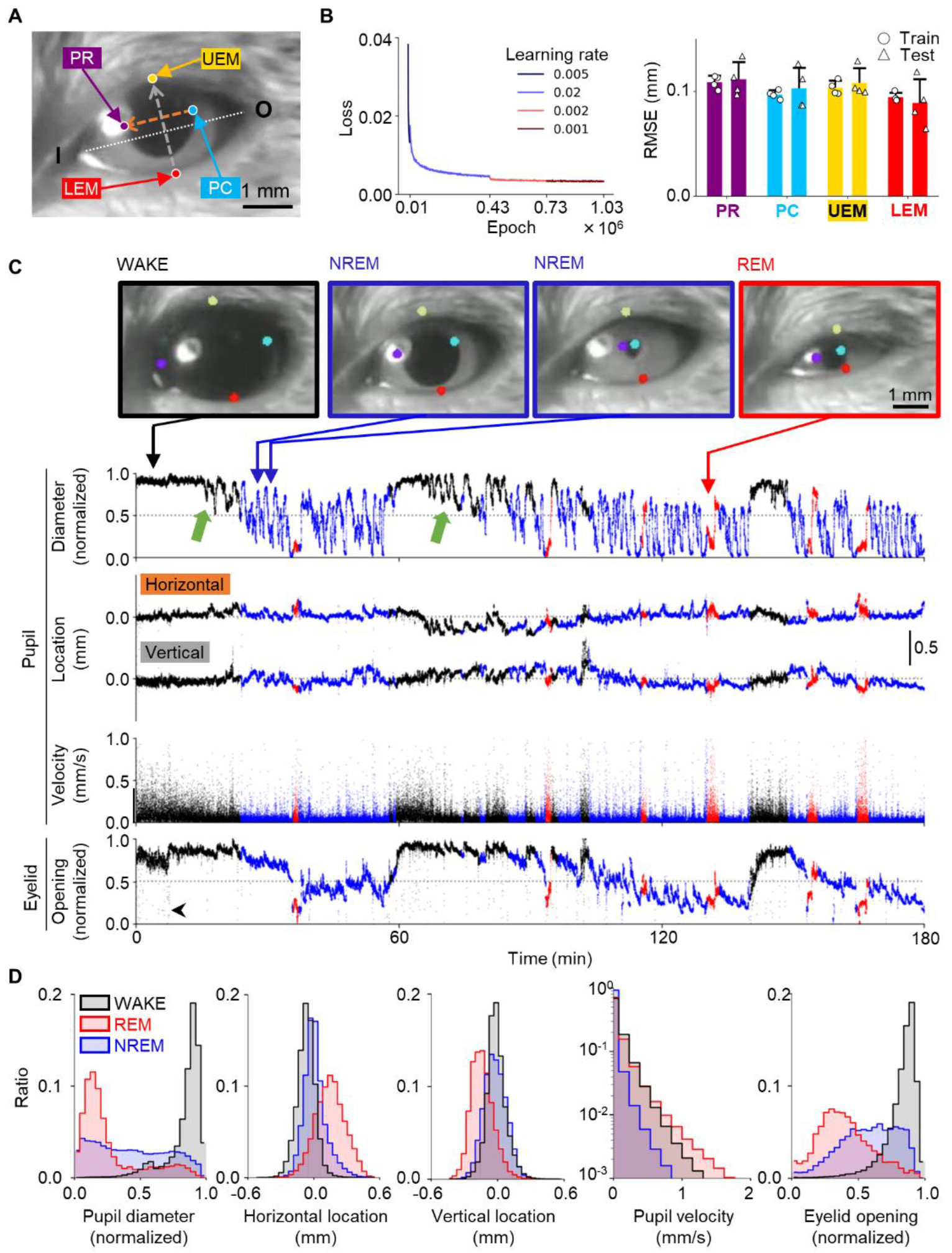
Preparation of pupil dynamics time-series as input feature variables for deep learning. **A:** Definition for manual positioning of pupil rostral (PR, purple dot), pupil caudal (PC, light blue dot), upper eyelid margin (UEM, yellow dot), and lower eyelid margin (LEM, red dot). Arrows in the orange and grey dashed line represent positive direction. I, O: inner and outer corners of the eye. **B:** Training and assessment of DeepLabCut for automatic positioning of PR, PC, UEM, and LEM in the left eye images. (Left) Learning curve of a cross-entropy loss (Loss) showed convergence of DeepLabCut training through a million epochs. Varying learning rates (distinct line colors) were used for optimal convergence. (Right) Generalization of the DeepLabCut estimation was confirmed by similar values of root mean square errors (RMSEs) for training data (open circles; 95% of the manually labeled 600 images) and test data (open triangles; 5% of the manually labeled images). Performance of DeepLabCut estimation deemed to be satisfactory because RMSEs were smaller than the minimum diameter of the mice pupil (*n* = 4 mice, 0.25 ± 0.02 mm). **C:** Video frame images of a mouse eye and time-series of pupillary dynamics from one recording session. (Top panels) Left eye images of a head-fixed mouse during WAKE, NREM, and REM episodes with colored dots denoting locations of rostral (purple) and caudal (light blue) edges of a pupil, and upper (yellow) and lower (red) limbs of an eyelid, which were obtained post-hoc using DeepLabCut (Supplementary Video). (Bottom traces) Time-series of pupil diameter (PD), horizontal and vertical pupil location (HL and VL), pupil velocity (PV), and eyelid opening (EO), which were calculated using the locations of the four-colored dots. In the pupil location panel, upward are the rostral and dorsal directions, respectively, for horizontal and vertical pupil location. Fluctuation in the pupil diameter occurred during NREM periods. We also noticed that fluctuation appeared during WAKE episodes that preceded NREM periods of approximately 10 min (green arrows). Eyelid opening sometimes showed a short deflection to a zero value in the time series (black arrowhead in the bottom trace), which corresponded to an eye blink. Traces were color-coded according to a supervisory hypnogram: WAKE, black; NREM, blue; REM, red. The temporal resolution of traces was 0.1 s. **D:** Histograms of pupil related feature variables during WAKE (black), NREM (blue), and REM (red) periods. Single feature variables at each time were not useful to determine sleep–wake states decisively because of their overlapping among vigilance states. For example, value of 0.9 for the pupil diameter appeared in all vigilance states.

Five features of pupil dynamics for pupil diameter (PD), pupil horizontal and vertical locations (HL and VL), pupil center velocity (PV), and eyelid opening (EO) were calculated based on time-series of the labeled positions for PR, PC, UEM, and LEM using a program that was custom written in Matlab (2020a; The MathWorks Inc., MA, USA). Specifically, values for the pupil dynamics time-series with estimated likelihood less than 0.99 by DeepLabCut were replaced with values obtained from linear interpolation of its adjacent values: 244,432 frames for PR (1.1%), 272,421 frames for PC (1.2%), 5,248 frames for UEM (0.023%), and 5,097 frames for LEM (0.023%) out of 22,589,862 images in all 39 sessions. After replacement, down-sampling to 10 Hz was applied. Then, PD was calculated as the distance between PR and PC, and scaled into the range of [0, 1] for each session. HL and VL were obtained as distances in millimeters from the average center position between PR and PC. Using the difference of the pupil position every 0.1 s, PV was calculated in millimeters per second. Also, EO was calculated as the distance between a UEM and an LEM, which were scaled into the range of [0, 1] for each session.

### Hypnograms

An EEG-based and EMG-based hypnogram was obtained using a conventional thresholding method for EEG and EMG amplitudes and a theta/delta power ratio with automatic correction.^7^ Specifically, the average power of the delta (1–4 Hz) and theta (6– 9 Hz) waves of the EEG, and of the EMG (30–300 Hz) over an entire session, were calculated. A 10-s bin was assigned as a WAKE period if the mean EMG power during the bin was greater than its session average. Otherwise, the bin was judged as NREM sleep if the delta wave power was larger than its session average and the theta/delta power was less than two.^45^ Alternatively, the bin was classified as REM sleep if the delta wave power was smaller than its session average and the theta/delta power was greater than two. The remaining bins which matched none of the criteria above were assigned a vigilance state of the preceding bin. Subsequently, automatic correction was applied to prohibit a single bin flipflop transition such as an NREM state to a Wake state then immediately back to an NREM state, which might represent a twitch behavior during sleep.^46,47^ Hereinafter, we designate hypnograms obtained using the automatic procedure described above as threshold-based hypnograms (Threshold in Fig. 3E). Each threshold-based hypnogram was then visually inspected and manually corrected to obtain the supervisory hypnogram for machine learning. Custom written Matlab codes were used for this vigilance state classification.

**Fig. 3.**
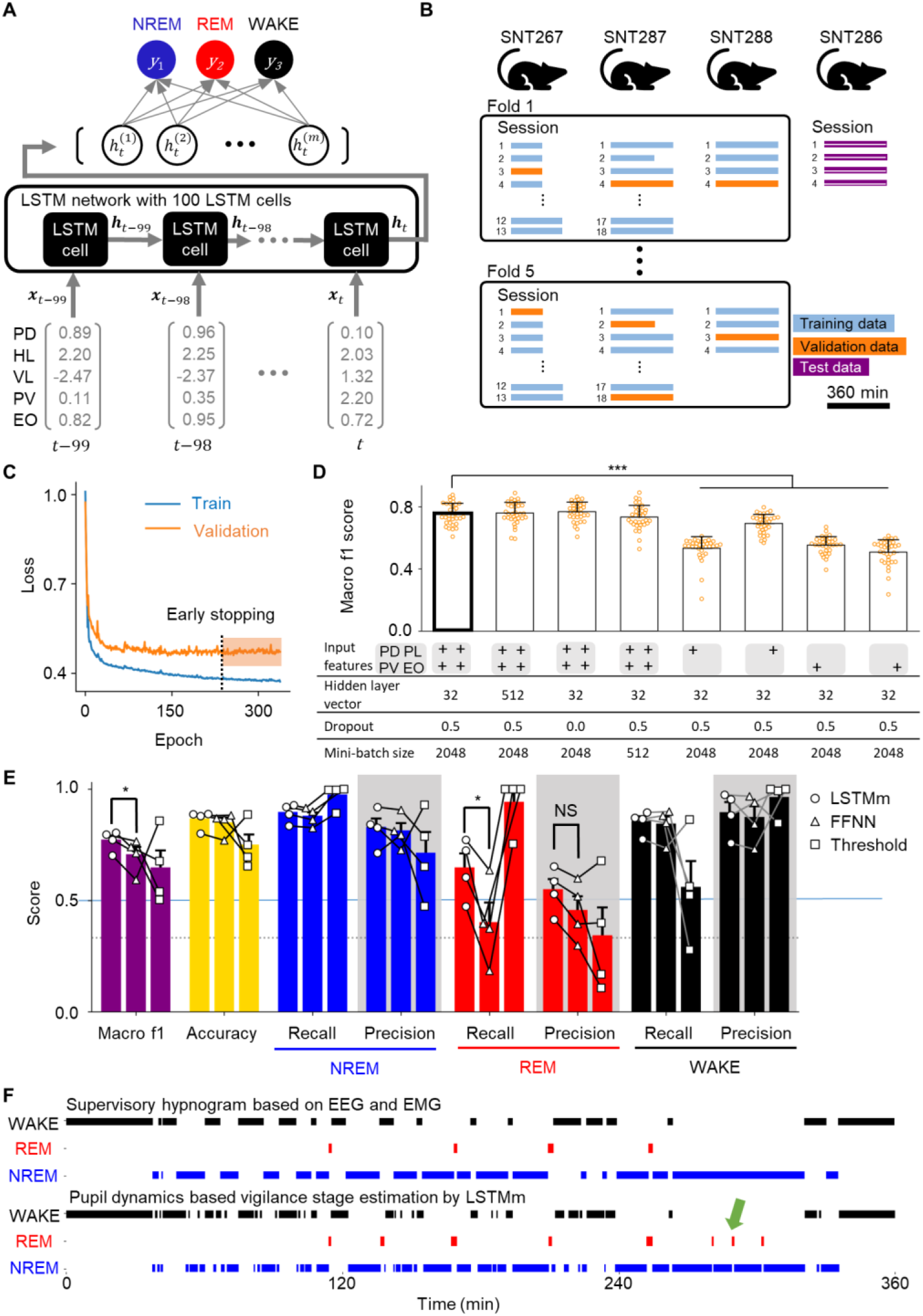
Pupil dynamics-based sleep-stage labeling using a deep neural network. **A:** Process flow of a long short-term memory (LSTM) network for pupil-based sleep stage classification. This example LSTM model had 5 input features for PD, HL, VL, PV, and EO. The feature vectors were sequentially processed in the LSTM cell to obtain a three-dimensional output vector, **y**, that consisted of log likelihood values for NREM, REM, and WAKE periods, the largest of which was used for estimation of the vigilance state at time *t*: PD, pupil diameter; HL and VL, pupillary horizontal and vertical locations; PV, pupil velocity; EO, eyelid opening. **B:** Assignment of datasets for training (blue), validation (orange), and testing (purple) of the LSTM networks to avoid data leakage. Stratified five-fold cross-validation technique was used for a hyperparameter tuning that compared performance of various LSTM architectures. Length of horizontal bars correspond to the session duration. **C:** Representative learning curves of an LSTM model in a single fold for hyperparameter optimization in fivefold cross-validation. The training loss (blue) approached a plateau, implying convergence of the model. Validation based early stopping was used to prevent overfitting of the network. In this example, training of the LSTM model was halted in the middle of learning at epoch 238 (vertical dotted line). **D:** The vigilance state classification performance of LSTM models was evaluated using a macro F1-score (worst 0, best 1) on validation data (orange bars in Fig. 3B) for hyperparameter optimization. We selected the LSTM model with the largest performance score as the hyperparameter optimized LSTM model (LSTMoptim; the leftmost thick bar) that exploited all input features (PD, PL, PV, and EO). The LSTMoptim performance was significantly higher than that of LSTM models with a single feature input (Dunnett’s test, *n* = 34 sessions, *** *P* < 0.001 vs. LSTMoptim as control). Pupil location (PL) includes features for pupillary horizontal and vertical locations (HL and VL). **E:** Comparison of the classification performance with the test data (purple bars in Fig. 3B) by the LSTMoptim with minor modification (LSTMm), the feed forward neural network (FFNN), and the conventional thresholding method without manual correction (Threshold). LSTMm and FFNN used pupil dynamics; Threshold used EEG and EMG for sleep staging. The overall classification performance was assessed based on Macro f1 and Accuracy. LSTMm showed higher overall performance than either FFNN or Threshold. Especially, LSTMm had a significantly higher macro F1-score than that of FFNN (purple bars for macro F1-scores: two-sided paired t-test: LSTMm 0.77 ± 0.02 vs. FFNN 0.70 ± 0.04, *\*p* = 0.027, *n* = 4 mice). The classification performance for each vigilance state was assessed with Recall and Precision (blue, red, and black bars). LSTMm had significantly higher Recall in REM than FFNN (red bars for Recall: two-sided paired *t*-test: LSTMm 0.64 ± 0.07 vs. FFNN 0.40 ± 0.09, \**p* = 0.0134, *n* = 4 mice). Note that higher REM Recall in Threshold was achieved at the expense of lower WAKE Recall. Statistical tests were adjusted for family-wise error using the Benjamini-Hochberg procedure with the false discovery rate of 0.05. **F:** Comparison of a supervisory hypnogram (top), and a hypnogram of LSTMm (bottom), demonstrating their overall mutual congruence. Characteristic differences between them were that NREM states in the supervisory hypnogram were estimated occasionally as REM in the LSTMm. Retrospective reexamination of EEG data during such periods indicated a transient theta power surge, implying that pupil dynamics might be a sensitive indicator of REM episodes.^48^

Pupil dynamics-based vigilance state classification of a head-fixed mouse was performed with a single layer long short-term memory (LSTM) neural network model (Fig. 3A). We compared performance of LSTM models with distinct hyperparameters including input features (PD, HL, VL, PV, EO) for 10 s at 10 Hz, batch sizes (512, 1024, 2048), hidden state vector dimensions (32, 128, 512), and dropout rates (0, 0.5) (Fig. 3D, **Table 1**). The feature vector input was processed sequentially for 100 time steps that corresponded to a 10-s bin. The first feature vector ***x***_*t*–99_ at time *t* – 99 was processed in the first LSTM cell to obtain an *m-* dimensional hidden state vector, ***h***_*t*–99_. This hidden state vector ***h***_*t*–99_ and the second feature vector ***x***_*t*–98_ were fed into the second LSTM cell to obtain the next hidden state vector: ***h***_*t*–98_. These processes were repeated 100 times to obtain the last hidden state vector, ***h**_t_*, which was then affine-transformed to produce a three-dimensional output vector, **y**, consisting of log likelihood values for NREM, REM, and WAKE episodes. The largest likelihood in the output vector was assigned as an estimated vigilance state at time *t*. We also prepared an FFNN model as reported earlier, but with a minor modification.^33^ The FFNN model consisted of an input layer for all five input features for 10 s duration, one hidden layer with 30 nodes, and a three-dimensional output layer for WAKE, NREM, and REM. The training procedure used for the FFNN model was the same as that used for the LSTM models.

**Table 1:** Evaluation results of hyperparameter tuning with 5-fold cross validation of LSTM.

| Hyperparameter |  |  |  |  | NREM |  | REM |  | WAKE |  |
| --- | --- | --- | --- | --- | --- | --- | --- | --- | --- | --- |
| input | batch size | hdim | dr | Macro f1 | Recall | Precision | Recall | Precision | Recall | Precision |
| PDPLPVEO | 2048 | 512 | 0.5 | $0.78 \pm 0.01$ | $0.83 \pm 0.03$ | $0.89 \pm 0.01$ | $0.64 \pm 0.02$ | $0.59 \pm 0.03$ | $0.9 \pm 0.02$ | $0.86 \pm 0.03$ |
| PDPLPVEO | 2048 | 512 | 0 | $0.78 \pm 0.01$ | $0.84 \pm 0.04$ | $0.88 \pm 0.01$ | $0.62 \pm 0.02$ | $0.61 \pm 0.02$ | $0.89 \pm 0.02$ | $0.86 \pm 0.03$ |
| PDPLPVEO | 2048 | 128 | 0.5 | $0.78 \pm 0.01$ | $0.83 \pm 0.04$ | $0.88 \pm 0.01$ | $0.65 \pm 0.02$ | $0.59 \pm 0.02$ | $0.89 \pm 0.02$ | $0.86 \pm 0.03$ |
| PDPLPVEO | 2048 | 128 | 0 | $0.78 \pm 0.01$ | $0.83 \pm 0.04$ | $0.88 \pm 0.01$ | $0.65 \pm 0.03$ | $0.57 \pm 0.03$ | $0.89 \pm 0.02$ | $0.86 \pm 0.03$ |
| **\*PDPLPVEO** | **2048** | **32** | **0.5** | **$0.78 \pm 0.01$** | **$0.84 \pm 0.04$** | **$0.88 \pm 0.01$** | **$0.63 \pm 0.02$** | **$0.6 \pm 0.03$** | **$0.89 \pm 0.02$** | **$0.87 \pm 0.03$** |
| PDPLPVEO | 2048 | 32 | 0 | $0.77 \pm 0.01$ | $0.83 \pm 0.04$ | $0.87 \pm 0.02$ | $0.63 \pm 0.04$ | $0.58 \pm 0.03$ | $0.89 \pm 0.02$ | $0.85 \pm 0.03$ |
| PDPLPVEO | 1024 | 512 | 0.5 | $0.78 \pm 0.01$ | $0.83 \pm 0.03$ | $0.88 \pm 0.01$ | $0.64 \pm 0.02$ | $0.59 \pm 0.03$ | $0.9 \pm 0.02$ | $0.86 \pm 0.03$ |
| PDPLPVEO | 1024 | 512 | 0 | $0.78 \pm 0.01$ | $0.83 \pm 0.03$ | $0.88 \pm 0.01$ | $0.62 \pm 0.02$ | $0.61 \pm 0.03$ | $0.9 \pm 0.01$ | $0.85 \pm 0.03$ |
| PDPLPVEO | 1024 | 128 | 0.5 | $0.78 \pm 0.01$ | $0.84 \pm 0.03$ | $0.88 \pm 0.01$ | $0.64 \pm 0.03$ | $0.59 \pm 0.03$ | $0.89 \pm 0.02$ | $0.86 \pm 0.03$ |
| PDPLPVEO | 1024 | 128 | 0 | $0.78 \pm 0.01$ | $0.83 \pm 0.03$ | $0.88 \pm 0.01$ | $0.64 \pm 0.02$ | $0.58 \pm 0.02$ | $0.9 \pm 0.02$ | $0.85 \pm 0.03$ |
| PDPLPVEO | 1024 | 32 | 0.5 | $0.78 \pm 0.01$ | $0.83 \pm 0.03$ | $0.88 \pm 0.01$ | $0.64 \pm 0.03$ | $0.59 \pm 0.03$ | $0.9 \pm 0.01$ | $0.86 \pm 0.03$ |
| PDPLPVEO | 1024 | 32 | 0 | $0.78 \pm 0.01$ | $0.83 \pm 0.03$ | $0.88 \pm 0.01$ | $0.64 \pm 0.02$ | $0.58 \pm 0.02$ | $0.9 \pm 0.02$ | $0.86 \pm 0.03$ |
| PDPLPVEO | 512 | 512 | 0.5 | $0.78 \pm 0.01$ | $0.84 \pm 0.04$ | $0.88 \pm 0.01$ | $0.64 \pm 0.03$ | $0.6 \pm 0.03$ | $0.89 \pm 0.02$ | $0.87 \pm 0.04$ |
| PDPLPVEO | 512 | 512 | 0 | $0.78 \pm 0.01$ | $0.83 \pm 0.04$ | $0.88 \pm 0.01$ | $0.65 \pm 0.03$ | $0.58 \pm 0.03$ | $0.89 \pm 0.02$ | $0.86 \pm 0.03$ |
| PDPLPVEO | 512 | 128 | 0.5 | $0.78 \pm 0.01$ | $0.83 \pm 0.03$ | $0.88 \pm 0.01$ | $0.63 \pm 0.03$ | $0.58 \pm 0.02$ | $0.9 \pm 0.01$ | $0.86 \pm 0.03$ |
| PDPLPVEO | 512 | 128 | 0 | $0.78 \pm 0.01$ | $0.83 \pm 0.03$ | $0.88 \pm 0.01$ | $0.63 \pm 0.02$ | $0.59 \pm 0.03$ | $0.9 \pm 0.01$ | $0.86 \pm 0.03$ |
| PDPLPVEO | 512 | 32 | 0.5 | $0.78 \pm 0.01$ | $0.83 \pm 0.03$ | $0.89 \pm 0.01$ | $0.64 \pm 0.03$ | $0.57 \pm 0.02$ | $0.91 \pm 0.01$ | $0.86 \pm 0.03$ |
| PDPLPVEO | 512 | 32 | 0 | $0.78 \pm 0.01$ | $0.83 \pm 0.04$ | $0.88 \pm 0.01$ | $0.64 \pm 0.02$ | $0.57 \pm 0.02$ | $0.9 \pm 0.02$ | $0.86 \pm 0.03$ |
| PD | 1024 | 512 | 0.5 | $0.67 \pm 0.04$ | $0.82 \pm 0.04$ | $0.82 \pm 0.02$ | $0.34 \pm 0.11$ | $0.36 \pm 0.1$ | $0.86 \pm 0.01$ | $0.84 \pm 0.03$ |
| PD | 1024 | 512 | 0 | $0.66 \pm 0.04$ | $0.82 \pm 0.02$ | $0.81 \pm 0.02$ | $0.32 \pm 0.11$ | $0.38 \pm 0.1$ | $0.85 \pm 0.02$ | $0.83 \pm 0.03$ |
| PD | 1024 | 128 | 0.5 | $0.66 \pm 0.02$ | $0.79 \pm 0.03$ | $0.81 \pm 0.02$ | $0.34 \pm 0.02$ | $0.39 \pm 0.05$ | $0.84 \pm 0.04$ | $0.82 \pm 0.03$ |
| PD | 1024 | 128 | 0 | $0.7 \pm 0.03$ | $0.81 \pm 0.04$ | $0.81 \pm 0.03$ | $0.46 \pm 0.04$ | $0.45 \pm 0.06$ | $0.84 \pm 0.03$ | $0.84 \pm 0.04$ |
| PD | 1024 | 32 | 0.5 | $0.55 \pm 0.01$ | $0.84 \pm 0.03$ | $0.79 \pm 0.02$ | $0.0 \pm 0.0$ | $0.0 \pm 0.0$ | $0.83 \pm 0.04$ | $0.83 \pm 0.03$ |
| PD | 1024 | 32 | 0 | $0.56 \pm 0.01$ | $0.83 \pm 0.04$ | $0.79 \pm 0.02$ | $0.03 \pm 0.03$ | $0.15 \pm 0.09$ | $0.83 \pm 0.03$ | $0.82 \pm 0.03$ |
| PD | 2048 | 512 | 0.5 | $0.63 \pm 0.03$ | $0.81 \pm 0.03$ | $0.81 \pm 0.01$ | $0.23 \pm 0.1$ | $0.31 \pm 0.09$ | $0.85 \pm 0.02$ | $0.83 \pm 0.03$ |
| PD | 2048 | 512 | 0 | $0.71 \pm 0.03$ | $0.8 \pm 0.03$ | $0.84 \pm 0.01$ | $0.44 \pm 0.04$ | $0.47 \pm 0.06$ | $0.87 \pm 0.02$ | $0.83 \pm 0.03$ |
| PD | 2048 | 128 | 0.5 | $0.65 \pm 0.01$ | $0.8 \pm 0.03$ | $0.79 \pm 0.02$ | $0.32 \pm 0.01$ | $0.37 \pm 0.04$ | $0.82 \pm 0.03$ | $0.82 \pm 0.03$ |
| PD | 2048 | 128 | 0 | $0.62 \pm 0.02$ | $0.81 \pm 0.03$ | $0.79 \pm 0.02$ | $0.22 \pm 0.08$ | $0.29 \pm 0.08$ | $0.83 \pm 0.03$ | $0.83 \pm 0.03$ |
| PD | 2048 | 32 | 0.5 | $0.56 \pm 0.02$ | $0.83 \pm 0.03$ | $0.79 \pm 0.02$ | $0.04 \pm 0.02$ | $0.13 \pm 0.09$ | $0.83 \pm 0.04$ | $0.82 \pm 0.03$ |
| PD | 2048 | 32 | 0 | $0.58 \pm 0.03$ | $0.84 \pm 0.03$ | $0.78 \pm 0.02$ | $0.08 \pm 0.06$ | $0.17 \pm 0.1$ | $0.81 \pm 0.03$ | $0.83 \pm 0.03$ |
| PD | 512 | 512 | 0.5 | $0.68 \pm 0.04$ | $0.79 \pm 0.04$ | $0.83 \pm 0.02$ | $0.38 \pm 0.1$ | $0.49 \pm 0.06$ | $0.87 \pm 0.02$ | $0.82 \pm 0.03$ |
| PD | 512 | 512 | 0 | $0.72 \pm 0.01$ | $0.81 \pm 0.03$ | $0.85 \pm 0.01$ | $0.49 \pm 0.05$ | $0.49 \pm 0.03$ | $0.89 \pm 0.01$ | $0.84 \pm 0.03$ |
| PD | 512 | 128 | 0.5 | $0.71 \pm 0.02$ | $0.8 \pm 0.03$ | $0.84 \pm 0.01$ | $0.45 \pm 0.05$ | $0.48 \pm 0.05$ | $0.88 \pm 0.01$ | $0.83 \pm 0.03$ |
| PD | 512 | 128 | 0 | $0.73 \pm 0.03$ | $0.81 \pm 0.04$ | $0.85 \pm 0.01$ | $0.5 \pm 0.05$ | $0.5 \pm 0.05$ | $0.87 \pm 0.02$ | $0.85 \pm 0.04$ |
| PD | 512 | 32 | 0.5 | $0.55 \pm 0.01$ | $0.85 \pm 0.03$ | $0.78 \pm 0.02$ | $0.0 \pm 0.0$ | $0.0 \pm 0.0$ | $0.82 \pm 0.04$ | $0.83 \pm 0.03$ |
| PD | 512 | 32 | 0 | $0.61 \pm 0.02$ | $0.82 \pm 0.04$ | $0.79 \pm 0.02$ | $0.22 \pm 0.07$ | $0.28 \pm 0.08$ | $0.81 \pm 0.04$ | $0.83 \pm 0.03$ |
| PL | 2048 | 32 | 0.5 | $0.72 \pm 0.01$ | $0.8 \pm 0.03$ | $0.83 \pm 0.02$ | $0.52 \pm 0.04$ | $0.53 \pm 0.04$ | $0.84 \pm 0.03$ | $0.81 \pm 0.03$ |
| PV | 2048 | 32 | 0.5 | $0.61 \pm 0.01$ | $0.8 \pm 0.03$ | $0.81 \pm 0.02$ | $0.19 \pm 0.01$ | $0.32 \pm 0.04$ | $0.79 \pm 0.04$ | $0.77 \pm 0.03$ |
| EO | 2048 | 32 | 0.5 | $0.51 \pm 0.02$ | $0.74 \pm 0.04$ | $0.76 \pm 0.02$ | $0.0 \pm 0.0$ | $0.1 \pm 0.1$ | $0.84 \pm 0.01$ | $0.75 \pm 0.04$ |
| PDPL | 2048 | 32 | 0.5 | $0.73 \pm 0.02$ | $0.81 \pm 0.03$ | $0.84 \pm 0.01$ | $0.51 \pm 0.06$ | $0.52 \pm 0.02$ | $0.87 \pm 0.02$ | $0.84 \pm 0.03$ |
| PDPV | 2048 | 32 | 0.5 | $0.77 \pm 0.01$ | $0.83 \pm 0.03$ | $0.87 \pm 0.01$ | $0.62 \pm 0.01$ | $0.58 \pm 0.02$ | $0.88 \pm 0.02$ | $0.86 \pm 0.03$ |
| PDEO | 2048 | 32 | 0.5 | $0.65 \pm 0.03$ | $0.79 \pm 0.03$ | $0.83 \pm 0.01$ | $0.28 \pm 0.09$ | $0.4 \pm 0.06$ | $0.88 \pm 0.01$ | $0.83 \pm 0.03$ |
| PLPV | 2048 | 32 | 0.5 | $0.74 \pm 0.02$ | $0.81 \pm 0.03$ | $0.85 \pm 0.02$ | $0.59 \pm 0.04$ | $0.54 \pm 0.03$ | $0.86 \pm 0.02$ | $0.83 \pm 0.03$ |
| PLEO | 2048 | 32 | 0.5 | $0.73 \pm 0.01$ | $0.8 \pm 0.03$ | $0.84 \pm 0.01$ | $0.56 \pm 0.04$ | $0.52 \pm 0.02$ | $0.87 \pm 0.01$ | $0.82 \pm 0.03$ |
| PVEO | 2048 | 32 | 0.5 | $0.72 \pm 0.02$ | $0.79 \pm 0.04$ | $0.86 \pm 0.01$ | $0.52 \pm 0.03$ | $0.49 \pm 0.04$ | $0.88 \pm 0.02$ | $0.82 \pm 0.04$ |
| PDPLPV | 2048 | 32 | 0.5 | $0.78 \pm 0.01$ | $0.83 \pm 0.04$ | $0.88 \pm 0.02$ | $0.63 \pm 0.03$ | $0.59 \pm 0.04$ | $0.89 \pm 0.03$ | $0.86 \pm 0.03$ |
| PDPLEO | 2048 | 32 | 0.5 | $0.74 \pm 0.01$ | $0.81 \pm 0.03$ | $0.86 \pm 0.01$ | $0.55 \pm 0.04$ | $0.54 \pm 0.04$ | $0.89 \pm 0.01$ | $0.84 \pm 0.03$ |
| PDPVEO | 2048 | 32 | 0.5 | $0.77 \pm 0.01$ | $0.82 \pm 0.03$ | $0.63 \pm 0.01$ | $0.90 \pm 0.01$ | $0.88 \pm 0.01$ | $0.58 \pm 0.02$ | $0.85 \pm 0.03$ |
| PLPVEO | 2048 | 32 | 0.5 | $0.75 \pm 0.02$ | $0.81 \pm 0.04$ | $0.87 \pm 0.02$ | $0.59 \pm 0.03$ | $0.55 \pm 0.03$ | $0.88 \pm 0.02$ | $0.83 \pm 0.03$ |

The LSTM architecture with the best performance (LSTMoptim) is shown in bold typeface with an asterisk: PD, pupil diameter; PL, pupil location; PV, pupil velocity; EO, eyelid opening; hdim, hidden state vector dimension; dr, dropout rate.

The entire dataset (*n* = 39 sessions, 4 mice) was assigned to training, validation, and testing of our neural network models (Fig. 3B). Data from a mouse SNT286 (4 sessions) were used only for the test data (purple bars in Fig. 3B) to prevent leakage of the test data into the training data. The remaining data (35 sessions from three mice: SNT267, SNT287, and SNT288) were used for training and evaluation of our neural network models with distinct hyperparameters using a stratified five-fold crossvalidation method. Specifically, the remaining 35 sessions were divided randomly into five groups, each including seven sessions. This session-by-session grouping was used to prevent leakages of information within a session. In a respective fold, four groups and the one remaining group were assigned respectively to training data (blue bars in Fig. 3B) and validation data (orange bars in Fig. 3B). These group assignments were repeated five times (5 folds) to cover all combinations of the five groups.

Training of neural networks was performed by backpropagation of the error algorithm using the adaptive moment estimation (ADAM) optimizer with hyperparameters: learning rate α = 7.5 × 10^−4^, gradient decay factor β_1_ = 0.9, and squared gradient decay factor β_2_ = 0.999. Learning curves of an LSTM network for each training step (epoch in Fig. 3C) were evaluated by an average weighted crossentropy loss in the training dataset (blue trace in Fig. 3C) and in the validation dataset (orange trace in Fig. 3C). The weights for WAKE, REM, and NREM were set as inversely proportional to bin counts for each vigilance state, avoiding underestimation of a minor class (REM) by assigning greater weight to its entropy loss. To avoid overfitting in a neural network and to avoid an increase of a validation loss, we used a validation-based early stopping when the validation loss did not decrease for more than 100 consecutive epochs (orange shaded rectangle in Fig. 3C) after training of at least 250 epochs, or when the training reached the maximum 500 epochs.

For hyperparameter optimization (Fig. 3D), the macro F1-score on the validation data (orange bars in Fig. 3B) was used to evaluate the vigilance state classification performance of various LSTM architectures. The macro F1-score, a harmonic mean of recall and precision, is a representative machinelearning metric for a multiclass classification with imbalanced data. The present dataset for vigilance state classification was moderately imbalanced because the minority class (REM) accounted for only a smaller proportion in % Total recording (3.5 ± 0.3%; Fig. 1E left panel), which might engender underestimation of REM episodes by the common metric accuracy. We designated the LSTM architecture with the highest macro F1-score as LSTMoptim (the leftmost thick bar in Fig. 3D). To achieve better performance, hypnography by the LSTMoptim was slightly modified, which we designated as LSTMm (Fig. 3E and 3F), such that if an estimated episode did not persist for three consecutive bins, i.e., ≥ 30 s, the estimated state was changed to the preceding episode. After hyperparameter optimization, the generalization performance for the vigilance states classification by LSTMm, FFNN, and Threshold was evaluated using test data (*n* = 4 sessions of SNT286; Fig. 3E). We used the metrices macro F1-score and accuracy to evaluate the vigilance states classification as a whole, and used the metrices recall and precision to evaluate the classification performance for each vigilance state. Neural network training was performed on Google Colaboratory pro using a Python library PyTorch (ver. 1.12.0). Data are expressed as mean ± SEM, except where noted otherwise.

### Data and code availability

Sample data for inputs (an eye movie and its timestamps), intermediate files (weights of LSTMoptim and DeepLabCut), and an output file (a pupil dynamics-derived hypnogram) are deposited in Mendeley Data (https://doi.org/10.17632/rr4gc6mybg.1). Python codes for LSTMoptim and DeepLabCut, and a Matlab code for pupil feature extraction are available online (https://github.com/gk-hazard/Pupil-to-Hypnogram).

## Results

### Preparation of a supervisory hypnogram based on EEG and EMG

EEG and EMG recordings and pupillometry were performed from a head-fixed mouse (Fig. 1A). A single recording session lasted 3–6 hr per day in a dark room during a light phase. Amplitudes of EEG and EMG from head-fixed mice during each vigilance state corresponded to that of freely behaving mice (Fig. 1B): The EEG amplitudes were larger during NREM than during either the WAKE or REM period. The EMG amplitudes were larger during WAKE than during either the NREM or REM period. We obtained first a hypnogram automatically using a conventional thresholding method for EEG and EMG, combined with automatic corrections to prohibit a single-bin flip-flop transition such as NREM-WAKE-NREM sequences. We designated this hypnogram without manual correction as a threshold-based hypnogram (Threshold). Then, a supervisory hypnogram was obtained by manual inspection and correction of the threshold-based hypnogram (Fig. 1C). The supervisory hypnogram for a head-fixed mouse reproduced the typical polyphasic nature of mouse sleep. Manual corrections of the vigilance states in the Threshold were necessary for 27 sessions among the 39 sessions from 4 mice (Fig. 1D).

Time spent in each vigilance state in a single recording session, and the duration for an individual episode in all recording sessions were calculated using a supervisory hypnogram from all data (Fig. 1E; total recording/session, 323 ± 63 min [mean ± SD]; 1672 episodes, 39 sessions, 4 mice; central mark in a box, median; edge of a box, 25^th^ and 75^th^ percentiles; whiskers, adjacent values). Actually, NREM sleep accounted for most of the total sleep time in a recording session (NREM, 48.9 ± 3.0%; REM, 3.5 ± 0.3%; NREM/[NREM + REM], 92.7 ± 0.5%), which was consistent with an earlier report on freely moving mice in light phase (NREM/[NREM + REM], ≈ 88 ± 5%).^2^ The median of episode durations (Fig. 1E, right panel) were 5.5, 1.7, and 3.0 min, respectively, for NREM, REM, and WAKE, which were also comparable to an earlier study for freely moving mice in the light phase (≈ 2.3, 1.5, and 1.1 min for NREM, REM, and WAKE).^33^

### Obtaining pupil dynamics time-series as feature inputs for a deep neural network

Pupillometry through wakefulness and sleep states was possible because head-fixed mice slept with incomplete closure of their eyelids (nocturnal lagophthalmos) as reported earlier.^33,35^ The reasons and mechanisms for open eyelids during sleep are unclear. Our mice slept with closed eyelids in their home cage, suggesting that the surgery did not prohibit closing of the eyelids.

We acquired positions of a pupil rostral (PR), a pupil caudal (PC), upper eyelid margin (UEM), and lower eyelid margin (LEM) in left eye movies using DeepLabCut (Fig. 2A). We prepared first supervisory data to train DeepLabCut by labeling the four positions manually in 600 left-eye images. The training of DeepLabCut was assessed using the cross-entropy loss (Loss) with 95% of the manually prepared dataset (570 from the 600 left eye images). Loss reached a plateau after a million epochs, indicating convergence of the training (Fig. 2B left). Generalization of the DeepLabCut position-estimation was evaluated using the root mean square error (RMSE) of the distance from manually defined supervisory positions to DeepLabCut-estimated positions. Acceptable generalization was achieved, as inferred from the similar RMSE found for the training data and test data (5% of the manually obtained dataset, i.e., 30 from the 600 left eye images), as shown in Fig. 2B (right, *n* = 4 mice, RMSE for training and test data: PR, 0.11 ± 0.01 mm, 0.11 ± 0.02 mm; PC: 0.10 ± 0.00 mm, 0.10 ± 0.02 mm; UEM, 0.10 ± 0.01 mm, 0.11 ± 0.01 mm; LEM, 0.09 ± 0.00 mm, 0.09 ± 0.02 mm). Indeed, no significant differences in RMSE was found between features (PR, PC, UEM, and LEM: F(3, 24) = 2.74, *p* = 0.07, two-way ANOVA), between datasets (training vs. test: F(1, 24) = 0.17, *p* = 0.69), or for their interaction (F(3, 24) = 0.28, *p* = 0.84). We concluded that the performance of our DeepLabCut estimation was satisfactory because the overall RMSEs for training data and test data (Fig. 2B right; *n* = 4 mice, 0.10 ± 0.01 mm, and 0.10 ± 0.01 mm) were smaller than the minimum diameter of the pupil (*n* = 4 mice, 0.25 ± 0.02 mm).

The trained DeepLabCut was used to estimate the four positions in the 22,589,940 left eye images in all movies automatically from 39 sessions of 4 mice (Fig. 2C upper panels, Supplementary Video). The five features of the eye dynamics were derived from the four positions (Fig. 2C traces). Pupil Diameter (PD) is a normalized distance between PR and PC (top trace in Fig. 2C). Although PD was large for most cases during WAKE periods, we found a decreasing trend and fluctuations of PD from 5–10 min before the transition to NREM sleep (green arrows in Fig. 2C top trace). During NREM states, low-frequency fluctuation of PD was evident, as reported previously.^33^ The PD fluctuation amplitude seemed to increase with the NREM state duration, implying that larger PD fluctuation accompanied deeper NREM sleep. During REM sleep, PD was small, but sometimes rapid dilation of PD occurred immediately before awakening. Horizontal and vertical pupil locations (HL, VL) were calculated as distances from the mean intermediated positions of PR and PC (second row from the top in Fig. 2C traces). Antiparallel movements of HL and VL were observed frequently during REM sleep, indicating anterior downward movements of the pupil. Also, pupil velocity (PV) was calculated for the intermediate position between PR and PC (third row from the top in Fig. 2C traces). Results show that PV was greater during WAKE and REM than during NREM periods. Although a small fraction, the highest pupil velocity was found during REM periods (Fig. 2D pupil velocity). Periodic increases of PV during NREM periods might coincide with PD fluctuation. The normalized distance between UEM and LEM is eyelid opening (EO; the bottom row of Fig. 2C traces). On average, EO was greatest during WAKE, followed by NREM, and the smallest during REM, which were similar to the pupil diameter but with distinct distribution during NREM periods, i.e., distributions for PD and EO during NREM periods were, respectively, uniform and peaked (Fig. 2D Eyelid opening). The figure shows that EO exhibited a gradual decrease during NREM, reaching minimum values during REM. Occasionally, EO showed a surge immediately before the transition from REM to WAKE episodes, which was similar to PD dynamics. The rapid deflection of EO to the minima during WAKE and NREM periods corresponded to eye blinks (black arrowhead in the bottom row of Fig. 2C traces). The features above (PD, HL, VL, PV, and EO) cannot determine the vigilance states individually because of their overlaps among WAKE, REM, and NREM episodes (Fig. 2D), e.g., PD showed almost uniform distribution during NREM sleep.

### Sleep stage estimation with pupil dynamics using a deep neural network

We developed LSTM models that are able to estimate vigilance states from pupil dynamics (Fig. 3A). Inputs for the model consisted of one to five features such as PD and EO. The inputs were 10 s long in a 10 Hz sampling rate, resulting in 100 time steps, each of which was processed sequentially with 100 LSTM cells. The hidden layer vector of the last LSTM cell, ***h**_t_*, was affine-transformed into a threedimensional output vector, **y**, that consisted of log likelihoods for NREM, REM, and WAKE states at time *t*. The state with the largest log likelihood was assigned as the estimated vigilance state of the model. Then, the estimation was compared to the supervisory hypnogram based on EEG and EMG to train the LSTM model. To avoid data leakage that might engender overestimation of the model performance, we 1) set aside a data of a mouse as test data (purple bars in Fig. 3B; 4 sessions of the mouse SNT286) from the entire dataset (39 sessions of 4 mice), and 2) performed hyperparameter optimization using a stratified five-fold crossvalidation method on a session-by-session basis (blue and orange bars in Fig. 3B). Consequently, we avoided data leakage in tests and validation because 1) a generalization performance of a model was evaluated with test data that were used neither for training nor validation, and because 2) session data that were used for training were not used for validation during hyperparameter optimization.

Supervised training of LSTM models was performed using validation-based early stopping to avoid overfitting (orange rectangle in Fig. 3C). After convergence in training, we compared the respective classification performance results obtained using LSTM models with distinct hyperparameters such as input features (distinct combinations of PD, PP, PV, and EO), a size of a hidden layer vector for each LSTM cell (***h**_t_*: 32, 138, or 512), a dropout rate (0 or 0.5) for the final hidden state ***h**_t_*, and a size for minibatch gradient descent (512, 1024, or 2048). The comparison was based on the average macro F1-scores using validation data over five-fold (Fig. 3D). The largest macro F1-score was achieved using the LSTM model with all input features, a hidden layer vector size of 32, a dropout rate of 0.5, and a minibatch size of 2048 (the leftmost thick bar in Fig. 3D; 0.78 ± 0.01 [mean ± SEM of 5 folds data]). The model with the highest macro F1-score was designated as the hyperparameter-optimized LSTM model (LSTMoptim). The score of LSTMoptim was significantly higher than those of LSTM models with a single input feature (right four bars in Fig. 3D; Dunnett’s test, *n* = 34 sessions, *** *P* < 0.001 vs. LSTMoptim as a control), although the performance score of LSTMoptim was not significantly different from those of other LSTM models with all input features (PD, HL, HV, and EO), irrespective of other hyperparameters for hidden layer unit sizes, dropout rates, or mini-batch sizes (left four bars in Fig. 3D). It is noteworthy that the LSTM model with a single feature for an eyelid opening (EO; the rightmost bar in Fig. 3D) showed comparable performance to that obtained using the model with a single feature for a pupil diameter (PD; the fourth bar from the right in Fig. 3D), implying that EO was as informative as PD, a commonly used feature for sleep–wake discrimination.^30^ **Table 1** presents the respective performances of LSTM models with distinct hyperparameters.

The generalization performance for vigilance state estimation was evaluated using the test data (Fig. 3E). To achieve better performance, the original hypnogram produced using LSTMoptim was modified slightly and automatically by introducing a criterion for vigilance state persistence. If an estimated vigilance state by LSTMoptim did not persist for at least three consecutive bins (≥ 30 s), then the estimated state was changed to the state immediately preceding it. We designated this modified hypnogram as LSTMm. The rationale for this modification was that we noticed a short-duration flip-flop transition between WAKE and NREM states in the original LSTMoptim during WAKE periods preceding NREM periods, which was apparently attributable to PD fluctuation immediately before the WAKE to NREM transition that resembled PD fluctuation during NREM periods (green arrows in Fig. 2C). This modification changed 4.6% of vigilance states by LSTMoptim (379 bins in 8736 bins in the test data), and significantly improved the scores for the macro F1-score, recall of REM episodes, and precision of REM episodes in LSTMm (*n* = 4 sessions, paired *t* test, family-wise error was corrected with FDR for *p* = 0.05, LSTMm vs. LSTMoptim: macro F1-score, 0.77 ± 0.02 vs. 0.73 ± 0.03, \**p* = 0.0093; REM recall, 0.64 ± 0.07 *vs* 0.57 ± 0.07, *p* = 0.11; REM precision, 0.55 ± 0.05 vs. 0.45 ± 0.05, \**p* = 0.0019).

We compared the performances of the LSTMm, the feed forward neural network (FFNN), and the threshold-based method (Threshold) (Fig. 3E). These methods require no manual correction of the hypnogram: for hypnogram estimation, LSTMm and FFNN used pupil dynamics; Threshold used EEG and EMG. Overall performance for vigilance state estimation was assessed with Macro f1 and Accuracy (purple and yellow bars in Fig. 3E). Results show that LSTMm performed better than either FFNN or Threshold. Particularly the macro F1-score of LSTMm was significantly higher than that of FFNN (purple bars in Fig. 3E, two-sided paired *t*-test: LSTMm 0.77 ± 0.02 vs. FFNN 0.70 ± 0.04, \**p* = 0.027, *n* = 4 mice). The classification performance for each vigilance state was assessed using Recall and Precision (blue, red, and black bars in Fig. 3E). LSTMm performed better than FFNN for all six comparisons. Particularly, LSTMm achieved a significantly higher recall for REM than FFNN (red bars in Fig. 3E, two-sided paired *t*-test: LSTMm 0.64 ± 0.07 vs. FFNN 0.40 ± 0.09, \**p* = 0.0134, *n* = 4 mice). The precision for REM were also higher in LSTMm than FFNN, although the difference was not significant (red bars in Fig. 3E, two-sided paired *t*-test: LSTM 0.55 ± 0.05 vs. FFNN 0.45 ± 0.07, *p* = 0.144, *n* = 4 mice). Although Threshold showed high recall for NREM and REM (0.97 ± 0.01 and 0.94 ± 0.02; blue and red bars for Threshold in Fig. 3E), these scores were achieved at the expense of lower scores for NREM and REM precision (0.71 ± 0.10 and 0.34 ± 0.04; blue and red bars for Threshold in Fig. 3E), and WAKE recall (0.56 ± 0.04; black bar for Threshold in Fig. 3E). Indeed, misidentification of WAKE states as NREM or REM states is apparent in Fig. 1D, with indications of higher Recall and lower Precision for NREM and REM (blue and red bars in Fig. 3E), and lower Recall and higher Precision for WAKE (black bars in Fig. 3E).

Time series of vigilance state labeling by LSTMm were compared with a supervisory hypnogram (Fig. 3F), demonstrating congruent estimation. We noticed more REM states in the hypnogram by LSTMm than in the supervisory hypnogram (green arrow in Fig. 3F). Retrospective inspection of EEG data during these REM episodes by LSTMm revealed a transient theta power increase in EEG during the 10-s bin that was labeled as an NREM state in the supervisory hypnogram, suggesting pupil dynamics as a sensitive indicator of REM, which was in accordance with the earlier report of a theta-pupil interaction.^48^

## Discussion

We demonstrated continual vigilance state estimation of a head-fixed mice based on 10 s pupil-dynamics using a single-layer LSTM model. The overall estimation performance would be satisfactory because the accuracy of sleep staging by our pupil dynamics-based estimator (86.4 ± 4.3%, mean ± SD) was comparable to the inter-rater agreement rate achieved by human experts (90–96%).^11,41,49^ Moreover, the performance approached that of recently reported EEG-based and EMG-based sophisticated neural networks such as a combination of a convolutional neural network (CNN) plus a hidden Markov model (93 ± 3.4% – 99 ± 0.4% for mice cohorts)^38^ and a CNN plus a bidirectional LSTM model (95.0–96.7%)^11^. The classification accuracy of these EEG-based and EMG-based estimators developed for freely moving mice might be lower when applied to head-fixed mice because the EMG activity of the head-fixed mice during WAKE periods seemed not to be consistently large enough to be clearly distinguishable from those during sleep periods. Indeed, the conventional Threshold method misassigned many WAKE periods as REM or NREM episodes (Fig. 1D), resulting in lower Recall for WAKE episodes (Fig. 3E). One advantage of the pupil dynamics-based sleep staging over an EEG-based and EMG-based staging is expected to be its compatibility with wide-field imaging of the cortex because pupil imaging requires no invasive surgery for electrode positioning on the thin skull of mice that would interfere with the imaging field.^50^

Great room remains for the improvement of pupil dynamics-based vigilance state estimation performance. The performance of FFNN and RNN are known to be similar if there are no strong temporal dependence on the data.^51,52^ Consequently, higher performance of our LSTM model than the FFNN model implies that the temporal dependence of pupil dynamics was important for sleep staging, even with 10-s time series. Therefore, the incorporation of pupil dynamics over a longer duration is expected to improve performance because the pupil diameter (PD) fluctuation preceded NREM sleep by approximately 10 min (green arrows in Fig. 3F). Moreover, the amplitude of PD fluctuation seemed to increase along with the duration of NREM episodes beyond approximately 10 min. Long-term pupil dynamics of such lengths are impossible to capture in the present LSTM model with feature vector inputs of 10 s. Indeed, characteristic mislabeling by our naïve LSTM model (LSTMoptim) did occur. For example, WAKE periods of approximately 10 min preceding NREM periods were occasionally estimated incorrectly as a repetition of WAKE and NREM states. The preceding PD fluctuation during WAKE periods might have been misrecognized by LSTMoptim as PD fluctuation during NREM sleep. This mislabeling was corrected in the proposed LSTM model (LSTMm) by introducing a simple rule of at least three consecutive vigilance states. Longer duration of feature vector inputs to a LSTM model might obviate the arbitrary rule and improve the scoring performance by constraining sleep staging from a prolonged perspective. Furthermore, improving the quality of feature vector inputs themselves is expected to improve the sleep staging performance by an LSTM model. For example, using a shorter bin duration, e.g., 4 s instead of the current 10 s, for the supervisory hypnogram can be expected to improve the sleep scoring performance because we noticed that a 10-s bin that had been mislabeled as a REM period by LSTMm during an NREM period in a supervisory hypnogram accompanied a transient increase of theta powers in EEG. This observation is consistent with an earlier report describing that a longer bin duration (20 s) for sleep – wake scoring of mice tended to miss very short events. Moreover, a higher frame rate video (> 60 Hz) can be expected to improve REM state detection because the frame rate of the current USB camera (approximately 33 Hz) is too slow for complete capture of rapid eye movement of mice recorded for 2–3 frames during movement.^33^

A few modifications would be necessary to make the current offline sleep-staging method online. Although the processing of LSTMm was completed rapidly and with only negligible resources, e.g., only a few seconds to process 6 hr feature vector inputs, preprocessing of pupil video images was time-consuming. The pre-trained DeepLabCut took 2.5 hr to process a 6 hr pupil movie that included > 700 million frames. The bottleneck was uploading the movie to the Google Colab website and doing DLC processing online. These findings indicate that a local computer for a DLC processing might be necessary for online sleep staging. Another issue for online sleep staging is that input features for pupil diameter and eyelid opening were relative values in this study, requiring whole session data to calculate them. Using absolute, instead of relative, values might resolve the issue. Applying the current method to a freely moving rodent would be a challenge because mice usually sleep with closed eyes. One possible solution for freely moving rodents might be a pupillometry through eyelids by combining a headmount infrared back-illumination pupillometry (iBip),^33^ a miniaturized ocular-videography system^53^ and scatter-resistant single photon imaging.^54^ Another limitation of the present method arises from its application to albino mice: we were unable to detect their pupils using a USB camera under infrared-red illumination.

Pupillary dynamics can be expected to provide fruitful opportunities to elucidate brain states of mice, rather than just providing an alternative to EEG-based and EMG-based hypnography.^55,56^ While training our LSTM model to classify vigilance states into three stages in the current study, we noted distinct pupillary dynamics even during a vigilance state that was assigned as a single state based on EEG and EMG. During WAKE periods, we noticed that pupil diameter fluctuation preceded the onset of NREM sleep which was difficult to capture with EEG and EMG. Extracellular spike recording has shown that mesopontine cholinergic neurons decreased their discharge frequencies 10–20 s before NREM sleep onset.^57^ Therefore, the preceding oscillations of pupil diameter may might imply the presence of neurons whose activity changes longer periods (several minutes) ahead of NREM sleep onset.^58,59^ In a human study, spontaneous waves of constriction and dilation of the pupils were attributed to decreased levels of wakefulness.^30^ Considering a recent finding that a hypnagogic period, which is a twilight zone between wakefulness and sleep, ignited creative sparks in human subjects,^60,61^ it is intriguing to speculate that mice pupillary oscillations preceding NREM sleep might be a useful measure to investigate sleepiness and creativity in mice. During NREM sleep, pupil diameter (PD) fluctuates with increasing amplitude along with NREM duration,^33^ indicating that mouse NREM sleep was not a monotonic state but was instead composed of different sleep depths, similar to that of humans.^62^ Recently, one report described that an oscillatory population-level activity of dorsal raphe serotonin neurons gradually increased its amplitude during NREM sleep,^63^ which implies a relation between deeper NREM sleep and larger PD fluctuations. During REM sleep, two microstates had been defined by eye movement frequencies.^64,65^ Phasic REM sleep accompanied bursts of eye movements, and tonic REM sleep had less eye movement, which are another example of a subdivision of a brain state suggested by pupil dynamics. Additionally, we noticed rapid dilation of the pupil diameter (mydriasis) at the end of REM sleep when a transition from REM to WAKE states occurred. This pupillary dilation might indicate neuronal activation of the noradrenergic locus coeruleus and related circuits in the brain.^66^ In conclusion, our simple and inexpensive approach of measuring pupil dynamics using a conventional USB camera would expand the repertoire for sleep stage scoring of head-fixed mice.

## Acknowledgments

This word was supported by JSPS KAKENHI Grants (Numbers 18H04952, 19KK0387 and 22H03033 to N.T.) and Keio University Academic Development Fund for Joint Research (N.T.).

**Supplementary Video. Pupil dynamics of a head-fixed mouse in a sleep-wake cycle.**

Left eye movies of a head-fixed mouse during WAKE, NREM, and REM periods for 10 s each. Positions for pupil rostral (PR, purple), pupil caudal (PC, light blue), upper eyelid margin (UEM, yellow), and lower eyelid margin (LEM, red) were obtained using DeepLabCut.

## Notes

https://doi.org/10.17632/rr4gc6mybg.1

https://github.com/gk-hazard/Pupil-to-Hypnogram

### Competing Interest Statement

The authors have declared no competing interest.

